# Single nuclei transcriptome of the Lesser Duckweed *Lemna minuta* reveals cell trajectories for an entire plant

**DOI:** 10.1101/2021.06.03.446947

**Authors:** Bradley W. Abramson, Mark Novotny, Nolan T. Hartwick, Kelly Colt, Brian D. Aevermann, Richard H. Scheuermann, Todd P. Michael

## Abstract

The ability to trace every cell in some model organisms has led to the fundamental understanding of development and cellular function. However, in plants the complexity of cell number, organ size and developmental times makes this a challenge even in the diminutive model plant *Arabidopsis thaliana*. Here we develop the Lesser Duckweed *Lemna minuta* as a model with a reduced body plan, small genome size and clonal growth pattern that enables simultaneous tracing of cells from the entire plant over the complete developmental cycle. We generated a chromosome-resolved genome for the 360 megabase genome and defined the growth trajectory of the entire plant with single nuclei RNA sequencing. The *L. minuta* gene complement represents a primarily non-redundant set with only the ancient *tau* whole genome duplication shared with all monocots, and paralog expansion as a result of tandem duplications related to phytoremediation. Thirteen distinct cell types representing meristem, the leaf-stem fusion called a frond, and root-like tissues were defined using gene orthology with single cell expression from model plants, gene ontology categories, and cell trajectory analysis. Dividing meristem cells give rise to two main branches of root-transition and mesophyll cells, which then give rise to terminally differentiated parenchyma, epidermal and root cells. Mesophyll tissues express high levels of elemental transport genes consistent with this tissue playing a role in *L. minuta* wastewater detoxification. The *L. minuta* genome and cell map provide a paradigm to decipher developmental genes and pathways for an entire plant.

**Sentence summary:** Genome and single nuclei transcriptome of the Lesser Duckweed *Lemna minuta* enables tracing of all developmental, transitional and terminal cells of an entire plant.

## Introduction

Single cell sequencing ushered in a new era of biology where it is possible to characterize cell types and function with unprecedented detail. In plants, this has resulted in detailed single cell RNA-seq (scRNA-seq) and scATAC-seq (Assay for Transposase-Accessible Chromatin) datasets primarily on different organ types from well-studied model organisms (Denyer et al., 2019; Ryu et al., 2019; Dorrity et al., 2020; Gujas et al., 2020; Liu et al., 2020a; Marand et al., 2020; Song et al., 2020; Liu et al., 2020b; Lopez-Anido et al., 2021; Xu et al., 2021). All plant studies to date start with protoplast isolation, which has the potential to miss some recalcitrant cell types and often requires correction to control for transcriptional changes elicited by lengthy enzyme treatment and centrifugation steps (Ryu et al., 2019). In addition, there are different single cell approaches that generally either result in assaying many cells with lower gene coverage (10X Genomics) or fewer cells with deeper gene coverage (SMARTseq2) (Yamawaki et al., 2021). For example, SMARTseq2 sequencing coupled with Fluorescence Activated Cell Sorting (FACS) of nuclei, or single nuclei RNA sequencing (snRNA-seq), provides deep transcriptome coverage of individual nuclei (Krishnaswami et al., 2016) while avoiding loss of cells or introducing issues associated with the protoplast isolation timing. The use of snRNA-seq in plants is nascent but can extend as a broader application for studying abiotic/biotic treatments and whole plant analysis.

Most single cell studies to date have been on specific plant organ tissue, such as the root, meristem or inflorescence, that encompass limited cell trajectories. Duckweed, aquatic plants in the family Lemnaceae, provide a unique opportunity to follow cells through all developmental time scales from one population due to its minimal morphology and clonal pattern of growth (Fig. 1). These basal monocots are some of the smallest and fastest growing plants on Earth, ranging in size from under a millimeter to centimeter, with some species doubling in under a day (Michael et al., 2020). There are five genera in the Lemnaceae with three having a flat leaf-like structure called a frond and root-like structures that may act as rudders or anchors (*Spirodela, Lemna* and *Landoltia*); the other two lack roots and are spherical or flat (*Wolffia* and *Wolffiella*). Before the model plant *Arabidopsis thaliana* rose to prominence, Duckweed were an important system for reductionist biology leading to fundamental understanding of chronobiology, flowering time, and phytohormone action (Hillman, 1976; Hillman and Culley, 1978).

**Figure 1.**
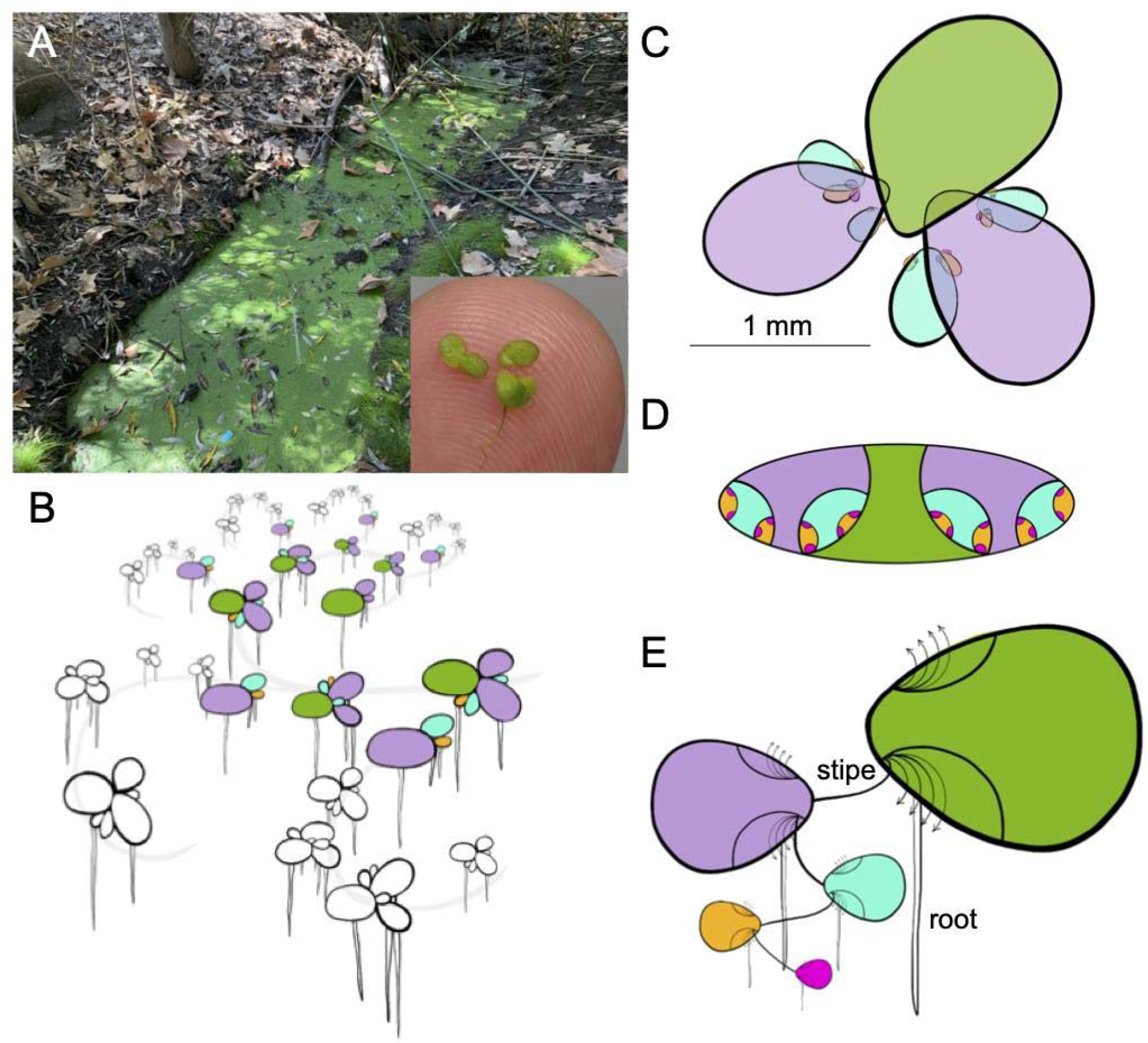
*Lemna minuta* growth highlighting its anatomical analogs to other plants. A) *L. minuta* (lm5633) growing as a dark green mat of fronds in a sewage slough at Cotton Creek Park, Encinitas, CA USA. The inset highlights the small size (~1 mm) of several fronds on a finger tip. B) Cartoon of the *L. minuta* generational contribution that leads to the dense frond mat: mother (M; green), daughter (D; purple), grand-daughter (GD; blue), great grand-daughter (GGD; orange) and great, great grand-daughter fronds (GGGD; pink). The grey line represents how they are connected through exponential growth. C) A single M frond view with the attached D fronds that have GD, GGD and GGGD fronds nested in the two meristematic pockets. D) A “nested doll” view of one M frond and the maturing generations of D, GD, GGD and GGGD in the pocket. E) One interpretation of the *Lemna* frond is that the M frond is an axillary stem that has two poaches or bracts where the D frond is attached by a stipe or internode. The D, GD, GGD and GGGD progression then is similar to a branching structure of a generic plant, and the root-like structure at each subsequent axillary node is equivalent to an adventitious root. The arrows indicate that multiple internodes will emerge from the axillary stem over time (Landolt, 1986).

Duckweed grow primarily through an asexual budding process where a mother frond gives rise to a daughter frond, forming a dense clonal population representing all stages of development in their lifecycle (Fig. 1, B-E). Despite being morphologically simple it has been hypothesized that the clonal population represents most complex tissues found in plants (Landolt, 1986). The mother front is an axillary shoot that gives rise to two pouches, or bracts, where the daughter frond is attached through a stipe, or internode, at the meristem and branches off in an alternating pattern similar to the growth of aerial tissue in more complex plants (Fig. 1E). Every daughter frond has 2 developing generations nested within pouches, which start to grow when the daughter is at the 18-cell stage and these new fronds start to differentiate at the 30-cell stage (Fig. 1D) (Rimon and Galun, 1968). Since the average lifespan of a duckweed plant is 30 days, and each frond is capable of generating 15-20 daughter fronds, the growth is exponential but also represents 256 (2^8^) fully formed but not expanded daughter fronds per mother frond (Fig. 1B). Therefore, a Duckweed population represents a three-dimensional look at a whole plant: developmental, cellular, and structural.

Duckweed also provide a compelling platform for genomic studies due to their core non-redundant gene set and relatively small genomes. The dawn of the genomics era yielded high quality genomes and transcriptomes for non-model systems, which resulted in Duckweed emerging once again as an attractive system to tackle problems of cell biology and development. *Spirodela polyrhiza*, The Greater Duckweed, was the first to be sequenced, revealing that in this 150 megabase (Mb) basal monocot genome there is a reduced complement of non-redundant protein coding genes (~19,000) representing a core plant proteome (Wang et al., 2014; Michael et al., 2017). Subsequently, genomes for *Lemna minor* (Van Hoeck et al., 2015), *Spirodela intermedia* (Hoang et al., 2020) and *Wolffia australiana* (Michael et al., 2020) have been published, revealing small genomes at 472, 160 and 375 Mb respectively. These sequenced genomes further support that Duckweed have a core plant gene set with few family expansions making them ideal for dissecting pathways and functional analysis.

Many species of Lemnaceae have dispersed widely beyond their natural ranges and are considered invasive species in their new habitats due to their rapid vegetative propagation (Moodley et al., 2016). A prime example of an alien invasive species is *Lemna minuta*, which is native throughout the temperate zones of the Americas, but has dispersed widely throughout Eurasia (Landolt, 1986). Analysis by Ceschin et al., maps the introduction of *L. minuta* in Europe to the 1950s -1960s with a dispersal rate of 40-50 km/per year (Ceschin et al., 2018). This dispersal process involved crossing of seas to reach places such as Ireland (Lucey, 2003) and Malta (Mifsud, 2010), which is consistent with a role for bird mediated dispersal, and hence the common name Duckweed (Silva et al., 2018). However, despite *L. minuta* doubling in roughly 24 hours, its growth rate does not explain its ability to outcompete native duckweed (*L. minor*) or other aquatic plants (Van Echelpoel et al., 2016; Paolacci et al., 2018).

The small organism size (~1mm), fast growth rate (~24 hrs), small genome size (~360 Mb), and ability to exploit diverse environments led us to select *L. minuta* as a model system to understand cell trajectories and function across an entire plant. *L. minuta* is emerging as a phytoremediation species due to its superior ability to remove various excess elements and toxins from wastewater (Ceschin et al., 2020; Fernández et al., 2020). This led us to hypothesize that the invasiveness of *L. minuta* may result from a specialized set of genes or cellular function, as it is apparently not solely due to its rapid growth rate. Therefore, we collected *L. minuta* from local wastewater (Fig. 1A) in order to ensure we captured the natural diversity of a successful clone. Duckweed clones sequenced to date have all been from the Landolt Collection, which have been maintained under aseptic lab conditions for 20-50 years (~1000s of generations). We brought a single sterile clone into culture (lm5633) and generated a chromosome-resolved reference genome. Additionally, snRNA-seq was performed on a population of clonally propagating plants to understand gene expression profiles across the whole plant’s cellular developmental landscape. The genome and snRNA-seq of lm5633 suggest mechanisms of invasiveness.

## Results

### Lemna minuta chromosome-resolved genome

*Lemna minuta* was collected from a wastewater run off in Cotton Creek Park, Encinitas, CA USA (33°2′58″N 117°17′29″W), sterilized and one clone was selected as a representative of the population for bulking (assigned the name lm5633 in the RDSC collection) (Fig. 1A). We estimated the genome size as 310 megabases (Mb) by k-mer (K = 31) frequency analysis using Illumina short reads sequencing (Fig. 2A), which was smaller than the reported average genome size estimated by flow cytometry of 365 Mb (Table 1) (Bog et al., 2020). The genome is highly heterozygous at 2.1% with 57% of the genome in high copy number elements (young transposable elements, centromeres, telomeres, rDNA arrays), while 28% is predicted to be repeat sequence (Fig. 2A; Table 1). The high level of heterozygosity suggests lm5633 may reproduce through outcrossing in the environment. We sequenced the genome using long-read Oxford Nanopore Technologies (ONT), and assembled reads into contigs using both our minimap/miniasm pipeline (Michael and VanBuren, 2020), as well as FlyE, the later of which produced a more contiguous assembly with a total length of 393 Mb, longest contig of 1.2 Mb and an N50 of 205 kb (Table 1).

**Figure 2.**
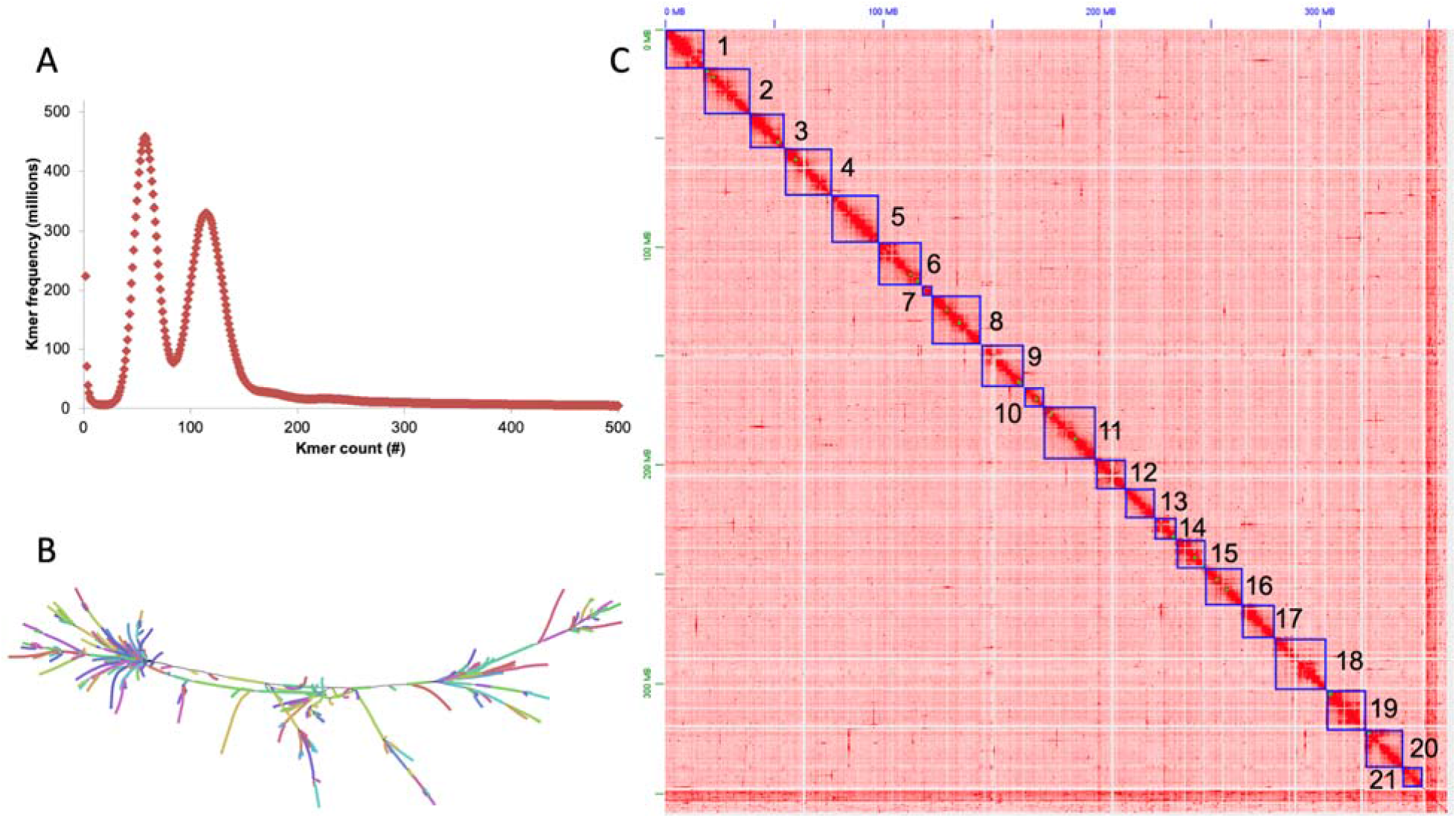
The highly heterozygous *Lemna minuta* (lm5633) genome resolved into 21 chromosomes. A) K-mer (k=31) frequency plot for lm5633 reveals two peaks consistent with a high level of heterozygosity (2%). B) Assembly graph of a 40 Mb region visualized with Bandage shows both the heterozygous branches as well as repeat tangles, “hairballs.” C) High throughput chromatin conformation capture (HiC) contact map resolving the polished lm5633 assembly into 21 chromosomes (darker red more contacts, lighter red to white less contacts).

**Table 1.** *Lemna minuta* (lm5633) genome statistics.

| **Table 1. *Lemna minuta* (Im5633) genome statistics.** |  |
| --- | --- |
|  | Im5633 |
| Estimated genome size flow cytometry (bp)* | 365,000,000 |
| Estimated genome size K-mer = 31 (bp) | 310,069,889 |
| Heterozygosity prediction K-mer31 (%) | 2.1 |
| High copy repeat sequence K-mer31 (%) | 57 |
| Chromosome-resolved genome size (bp) | 360,454,868 |
| Assembled genome size (bp) | 392,702,877 |
| Contig (#) | 9,473 |
| Longest contig (bp) | 1,268,926 |
| Contig N50 length (bp) | 205,076 |
| Scaffold (#) | 2,407 |
| Longest scaffold (bp) | 24,571,886 |
| Scaffold N50 length (bp) | 19,492,010 |
| Chromosomes after HiC (#) | 21 |
| BUSCO<sup>&</sup> complete (%) | 72.5 |
| Repeat predictions (%) | 58.2 |
| Telomere length average (bp) | 10,085 |
| Telomere length, longest (bp) | 25,271 |
| Predicted genes (#) | 22,873 |

\*Averaged over all *L. minuta* clones tested by (Bog et al., 2020)
<sup>&</sup>BUSCO library: liliopsida\_odb10

Since the genome assembly was larger than the predicted genome size we reasoned that some of the genome was retained in two haplotypes. The genome assembly graph confirmed that excess haplotypes were retained appearing as branches in the graph, as well as suggesting a complex repeat structure (transposable elements; TEs) resulting in repeat tangles, or “hairballs” in the graph (Fig. 2B). We assessed the completeness of the assembly with benchmarking universal single-copy orthologs (BUSCO) finding 72.5% complete (Table 1); the lower percent complete is consistent with other high quality chromosome resolved Duckweed genomes (Hoang et al., 2018; Harkess et al., 2020; Michael et al., 2020). In addition, we found 11.8% duplicated BUSCO genes consistent with the presence of additional haplotypes in the assembly (Table 1; Supplemental Table S1). We purged the excess overlapping haplotypes and scaffolded the contigs with Illumina-based high throughput chromatin conformation capture (HiC) that resolved the genome into 21 chromosomes consistent with published cytology (Fig. 2C) (Landolt, 1986). The final lm5633 HiC assembly was collinear with the high-quality assembly of *S. polyrhiza* clone 9509 (sp9509) chromosome structure (Fig. 3A).

**Figure 3.**
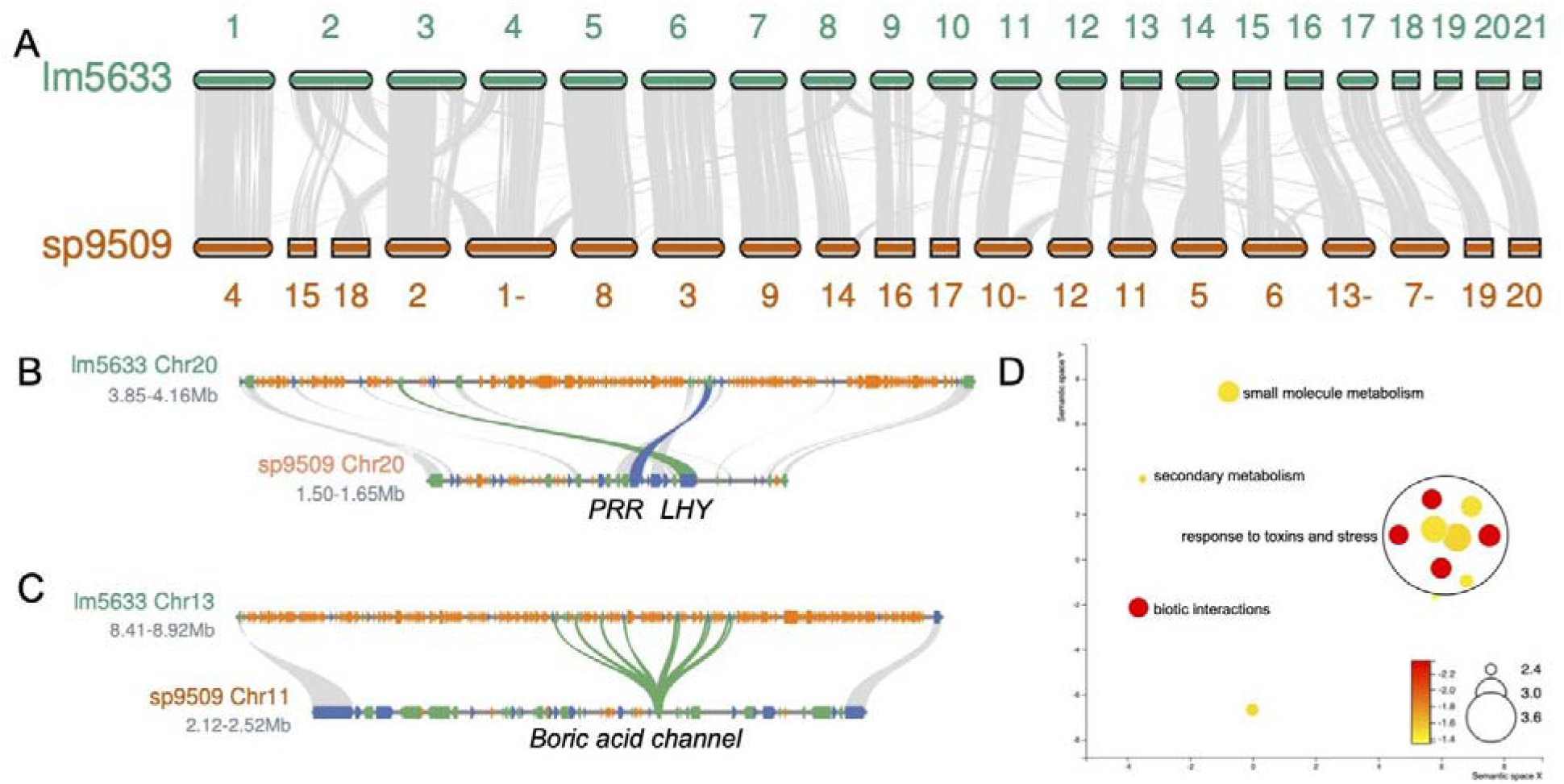
Gene family expansion in lm5633 is driven by tandem duplication (TD). A) Lm5633 chromosomes aligned to sp9509 chromosomes with syntenic blocks (grey lines) anchoring positions between the two genomes. Chromosomes are the correct ratio between one another but are not to scale between the two species. A minus sign after the number means the chromosome has been flipped for visualization purposes. B) The single MYB transcription factor *LATE ELONGATED HYPOCOTYL* (*LHY*; blue line) and *PSEUDO-RESPONSE REGULATOR 7* (*PRR7*; green line) are in tight linkage and TE fragments (orange) resulting in the region expansion in lm5633. Grey lines connect other syntenic genes (blue, forward; green reverse). C) Boric acid channel TD (green line) also showing the expansion of the lm5633 genome due to TE fragments (orange). D) Sympantic principle component analysis (PCA) of significant GO terms associated with TDs in lm5633. Size of the circle is the log frequency and the color (red high, and yellow low) is the log FDR.

### Lm5633 gene prediction and annotation

Long read assemblies cover more repeat sequences and usually allow the identification of putative centromere sequences, definition of telomere lengths, and annotation of full-length TEs (Michael and VanBuren, 2020). Consistent with the high copy number repeat k-mer frequency estimate, we identified that 58.2% of the genome was repeat sequence, which is double that of sp9509 and similar to *W. australiana* clone 8730 (wa8730) (Table 1; Supplemental Table S2). The lm5633 genome has a gypsy/copia ratio of ~2, similar to that found in sp9509, but distinct to wa8730 where the ratio is closer to 1 (Supplemental Table S2) (Michael et al., 2017; Michael et al., 2020). The larger genome size of lm5633 compared to sp9509 was primarily due to the increase in retained TE fragments, which is seen in the number of repeats found intervening in the evolutionarily conserved linkage of core circadian clock genes (Fig. 3B) (Michael et al., 2020). Similar to *S. polyrhiza*, we did not detect high-copy number centromere repeats (Michael et al., 2017), but we did identify telomere (AAACCCT) arrays that are longer (average=10 kb; longest 25 kb) than sp9509 (average=3 kb; longest=6 kb), yet shorter than what we have found in wa8730 (average=18 kb; longest=70 kb) (Supplemental Table S3) (Michael et al., 2020).

The sequenced duckweed genomes have the fewest protein coding genes found in angiosperms to date with wa8730 and sp9509 having just 14,324 and 18,507 genes respectively (Michael et al., 2017; Michael et al., 2020). After masking the repeat sequence, we predicted 22,873 protein coding genes in the lm5633 genome (Table 1), which is similar to the 22,382 and 22,245 protein coding genes predicted in the lm5500 and si7747 assemblies, respectively (Van Hoeck et al., 2015; Hoang and Schubert, 2017). The higher gene counts for both lm5633 and si7747 compared to wa8730 and sp9509 are a result of those genomes having orthogroups with greater than 10 genes, which is similar to *Arabidopsis* and rice where gene families are much larger (Fig. 4A). Of the 99 lm5633 orthogroups that had greater than 10 genes, 50% had between 0 and 2 genes in sp9509. The expanded lm5633 families could result from whole-genome duplication (WGD), tandem duplication (TD), proximal duplication (PD), transposed duplication (TRD), or dispersed duplication (DSD) (Qiao et al., 2019). The expanded lm5633 genes were predominantly a result of TD and PD events, as compared to sp9509, leading to expansion of genes involved in pathogen defense, response to stress, and nutrient acquisition (Fig. 3D; Supplemental Table S4). For instance, there is an 11 gene TD of a boric acid channel (Fig. 3C), which may reflect the ability of lm5633 to extract large amounts of boron from its environment, improving survivability, and invasiveness (Fernández et al., 2020).

**Figure 4.**
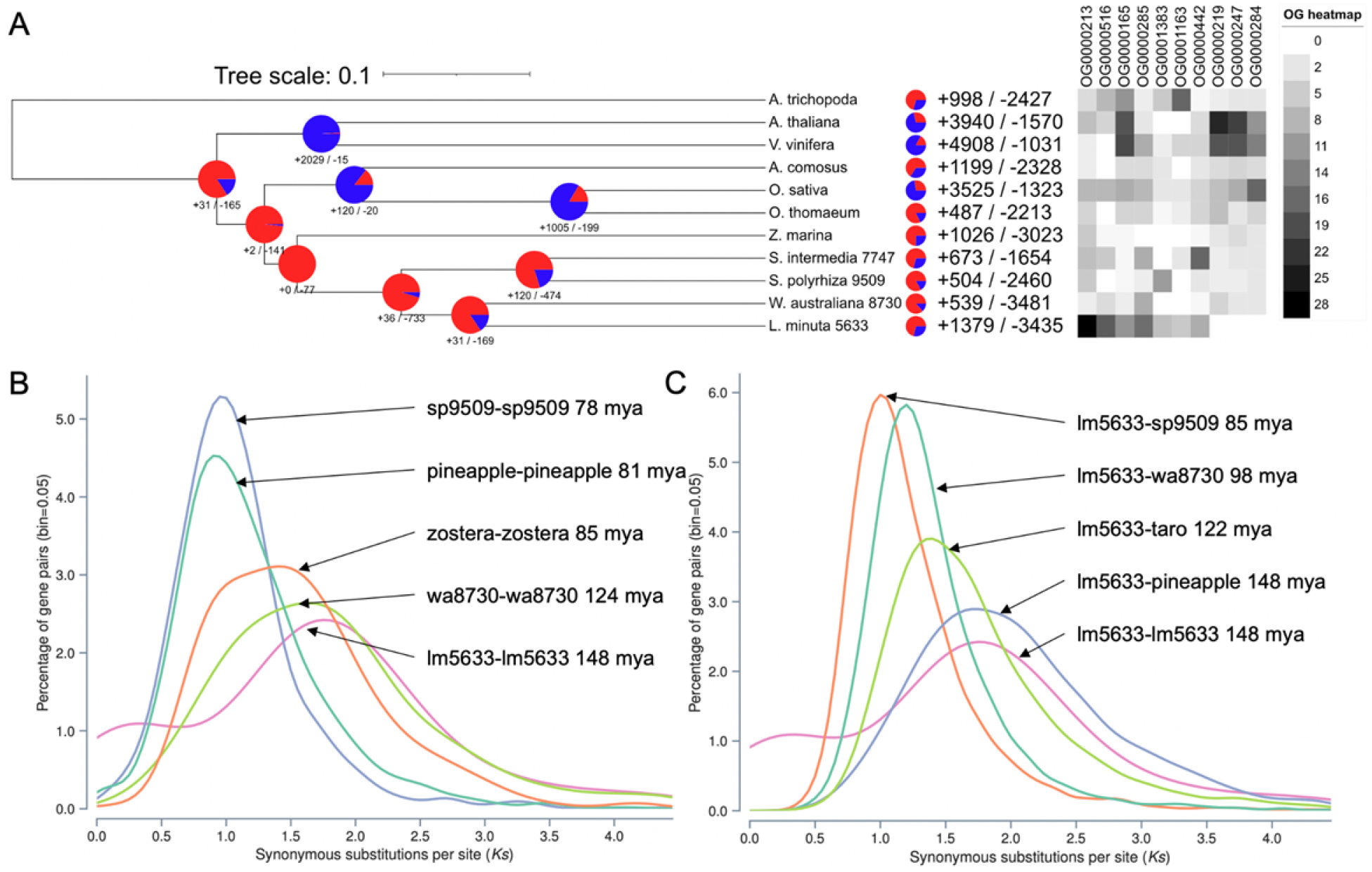
Whole genome evolution shows consistent gene family contractions in lm5633 ancestry. A) Gene family contractions (red) and expansions (blue) along the phylogenetic tree leading to minimized duckweed genomes are shown at each node and total protein family contractions and expansions for each species (right). Lm5633 shows the greatest similarity and gene family conservation with wa8730. A heatmap of orthogroups significantly expanded or contracted in lm5633. B) Self versus self and C) lm5633 versus other synonymous substitution (Ks) plots to elucidate the WGD history of lm5633. Dating is based on the mean Ks peak for each comparison using all paralogs/orthologs. Plant species used: *A. trichopoda* (Amborella), *A. thaliana, V. vinifera* (grape), *A. comosus* (pineapple), *O. sativa* (rice), *O. thomaeum* (oropetium), *Z. marina* (zostera), *C. esculenta* (taro), *S. intermedia* (si7747), *S. polyrhiza* (sp9509), *W. australiana* (wa8730), and *L. minuta* (lm5633).

A comparison of orthologous proteins between lm5633 and wa8730 revealed a number of expanding orthogroups relating to the ability of lm5633 to thrive in diverse environments. We found 14 significantly expanded orthogroups in lm5633 compared to the most recent common ancestor wa8730 (Fig. 4A). There are five orthogroups annotated with gene ontology (GO) terms: OG0000813, vacuolar membrane; OG0000014, signaling; OG0001260, defense response to fungus; OG0000015, detection of bacterium; OG0003377, delta14-sterol reductase activity. There are 3 significantly contracted orthogroups compared to wa8730 with one having an annotation for GO:0005787,signal peptidase complex. An additional expanded orthogroup, OG0000243, encodes a family of proteins relating to multidrug and toxic compound extrusion (MATE) proteins that have more than doubled from 2 copies in wa8730 to 5 copies in lm5633. The MATE transporters are often associated with increased plant resilience to toxic compounds and adaptability to metals including iron homeostasis (Upadhyay et al., 2019). These evolving orthogroups are consistent with the ability of lm5633 to thrive under adverse conditions of abiotic and biotic stress.

### Lm5633 *chromosome evolution*

The lm5633 genome was resolved into 21 chromosomes (Fig. 2C and 3A), which is consistent with cytological studies (2n=42) (Landolt, 1986); this is one more chromosome than sp9509 and three more than si7747 (Michael et al., 2017; Hoang et al., 2020). Since *S. polyrhiza* is thought to be the basal duckweed, we looked specifically at the synteny between the lm5633 and sp9509 chromosomes and found 18 chromosomes share high sequence collinearity with at least 5 that are completely collinear (Fig. 3A). There are several lm5633 chromosomes (2,19, and 21) that are the result of fragments from several sp9509 chromosomes, whereas no sp9509 chromosomes resulted from lm5633 fusions, consistent with the basal nature of the *S. polyrhiza* genome.

*Spirodela polyrhiza* has experienced at least two lineage specific WGDs (B”/a”) in the non-grass monocots (Wang et al., 2014; Ming et al., 2015). Consistent with this WGD history, sp9509 has a 3:1 syntenic depth compared to the *Amborella trichopoda*, which is the progenitor basal angiosperm without a WGD (Amborella Genome Project, 2013). In contrast, lm5633 has a 2:1 syntenic depth compared to *A. trichopoda*, suggesting lm5633 may have a higher level of fractionation (WGD followed by diploidization and gene loss) compared to sp9509. Consistent with lm5633 having a higher level of fractionation, only 4.1% of its paralogs remain in syntenic blocks compared to 49% in sp9509. Moreover, lm5633 and sp9509 have a 1:1 syntenic depth, sharing 88% and 86% gene pairs respectively, indicating that despite the higher level of fractionation in lm5633 the gene content from their shared WGD history is preserved (Fig. 3).

An alternative hypothesis explaining the lower level of paralogs in syntenic blocks in lm5633 is that it did not experience the *S. polyrhiza* B”/a” WGDs, and instead only experienced the *tau* (□) WGD that is shared across most of the monocot lineage (Jiao et al., 2014). Therefore, we looked at the synonymous substitution (Ks) rates of lm5633 paralogs to estimate their age, and found that both wa8730 and lm5633 lack a Ks peak corresponding to B”/a” WGD with only a peak corresponding the to the *tau* (□) WGD (Fig. 4B). Consistent with this, the divergence of lm5633 and pineapple coincides with the *tau* (□) WGD (Fig. 4C). These results suggest that the *Wolffia* and *Lemna* genera may not have the lineage specific WGDs that are found in *Spirodela*. Additional genomes from *Lemna, Wolffia* and *Wolffiella* will be required to fully understand the WGD history of duckweed. However, the Ks plots are consistent with the genome innovation of lm5633 resulting from recent TDs in genes associated with abiotic and biotic stress, which provide clues as to its invasive abilities.

### Defining cell types in lm5633 with snRNA-seq

The unique life cycle and growth pattern of *L. minuta* plants from mother to daughter and grand-daughter fronds covers all cell developmental stages and types providing an opportunity to follow the developmental trajectory of all cells (Fig. 1). We wanted to identify as many cell types as possible across the diverse developmental states with high gene coverage per cell. Therefore, we carried out single nuclei RNA sequencing (snRNA-seq), which enabled capturing an exact moment in development through the immediate freezing of tissue and avoiding long protoplasting steps that can result in unsampled cell types and sample preparation artifacts (Denyer et al., 2019). We isolated individual nuclei from a population of whole lm5633 plants grown under standard laboratory conditions by FACS and then prepared libraries with SMARTseq2 to achieve high gene coverage per nuclei. After quality filtering, 49% of the annotated lm5633 genes were expressed across 269 nuclei.

One problem encountered with non-model plants is the lack of annotation. For instance, only six of the twenty most highly expressed genes for all cells in this dataset had annotations, five related to photosynthesis as expected for green tissues (Fig. 5A). Therefore, we leveraged a combination of cell trajectory analysis, gene ontology (GO) annotation, and orthology prediction to define single cell marker genes from model plants for describing cell types in lm5633. Expression based dimensionality reduction resulted in 13 distinct cell clusters with a total of 1,733 significantly differentially expressed genes between clusters (Fig. 5B). To initially define the 13 clusters we identified lm5633 orthologs relative to published single cell studies in *Arabidopsis, O. sativa* (rice) and *Zea mays* (corn) (Denyer et al., 2019; Satterlee et al., 2020; Xu et al., 2021) (Supplemental Table S5). Of the 1,733 marker genes found for the 13 clusters, we found 140 marker genes that did not appear in any orthogroup and on average each marker gene was in an orthogroup with 2.3 copies per species (Supplemental Table S6). An average of 134 marker genes per cluster were determined with a total of 446 marker genes annotated based on model organism orthology, averaging to 18 annotated marker genes per cluster (Fig. 5C; Supplemental Table S7).

**Figure 5.**
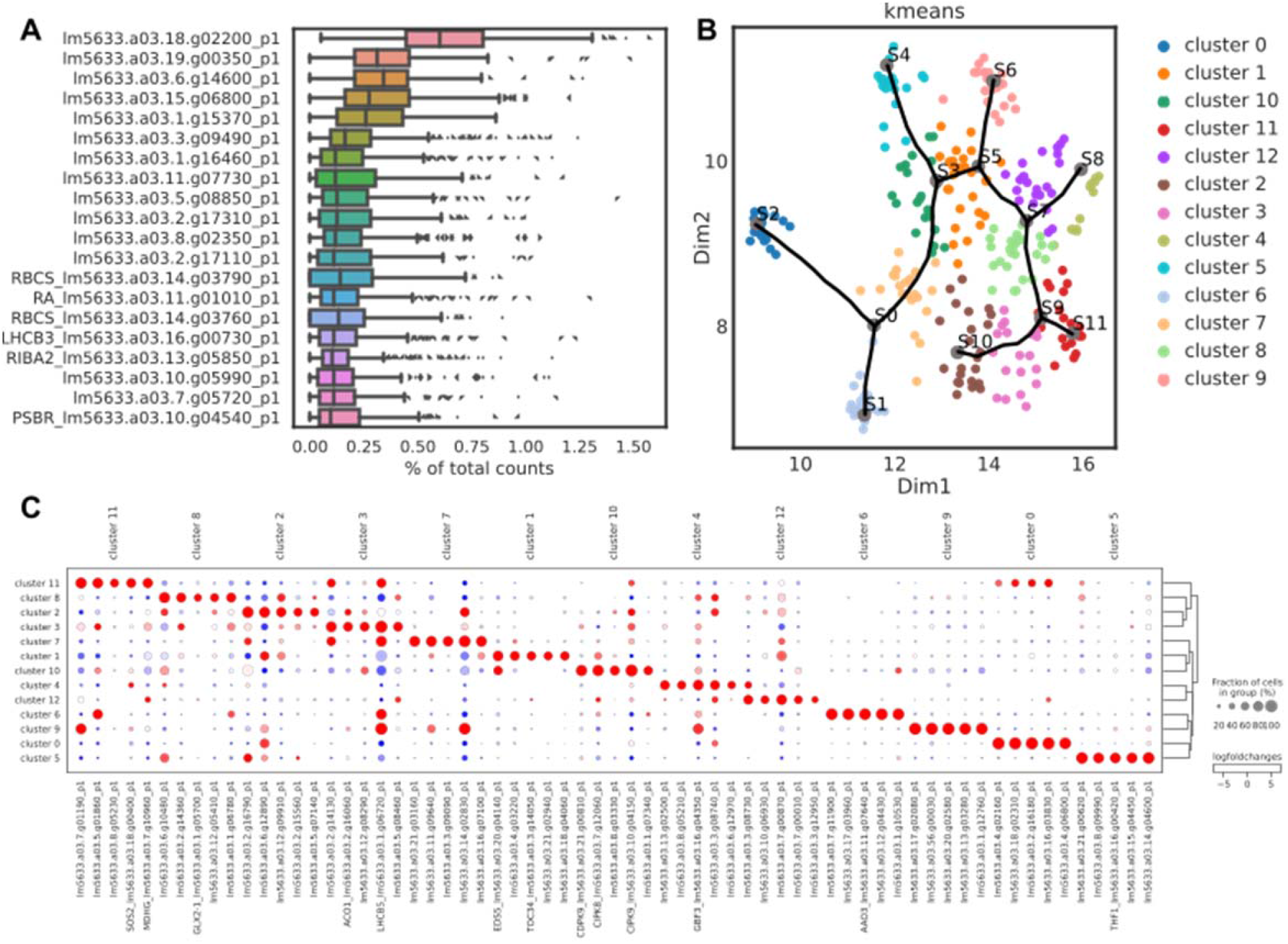
Single nuclei RNA sequencing (snRNA-seq) of clonally propagating whole lm5633 plants. A) The most abundant transcripts reveal that highly expressed genes are generally related to photosynthesis. B) UMAP embedding of 13 k-means clusters in the snRNA-seq dataset. Trajectory analysis (black line) reveals a complex network where multiple branches result in terminally fated cell types. C) Log fold change in marker genes for the 13 specific clusters shows robust separation of cell types for functional analysis. Expression level (red, high; blue, low; larger circle, more cells; smaller circles, fewer cells).

GO analysis of the marker genes provided additional functional support for the distinction of each cluster as well as potential functional overlap of cell types (Fig. 6A). Each cluster/cell type had unique GO terms while several had overlapping terms such as meristem-replication, meristem-axillary meristem, root epidermis-axillary meristem, root-axillary meristem, root transition I-axillary meristem, and meristem-root. These results suggest that the meristem-like tissue and different root cell types are intimately associated. Together with orthology from model plants (described below), the cell trajectory showed that meristem gives rise to root transition, meristem-like, and mesophyll cells; mesophyll then gives rise to parenchyma followed by epidermis, while root transitions gives rise to root epidermis cells followed by root cells (Fig. 7).

**Figure 6.**
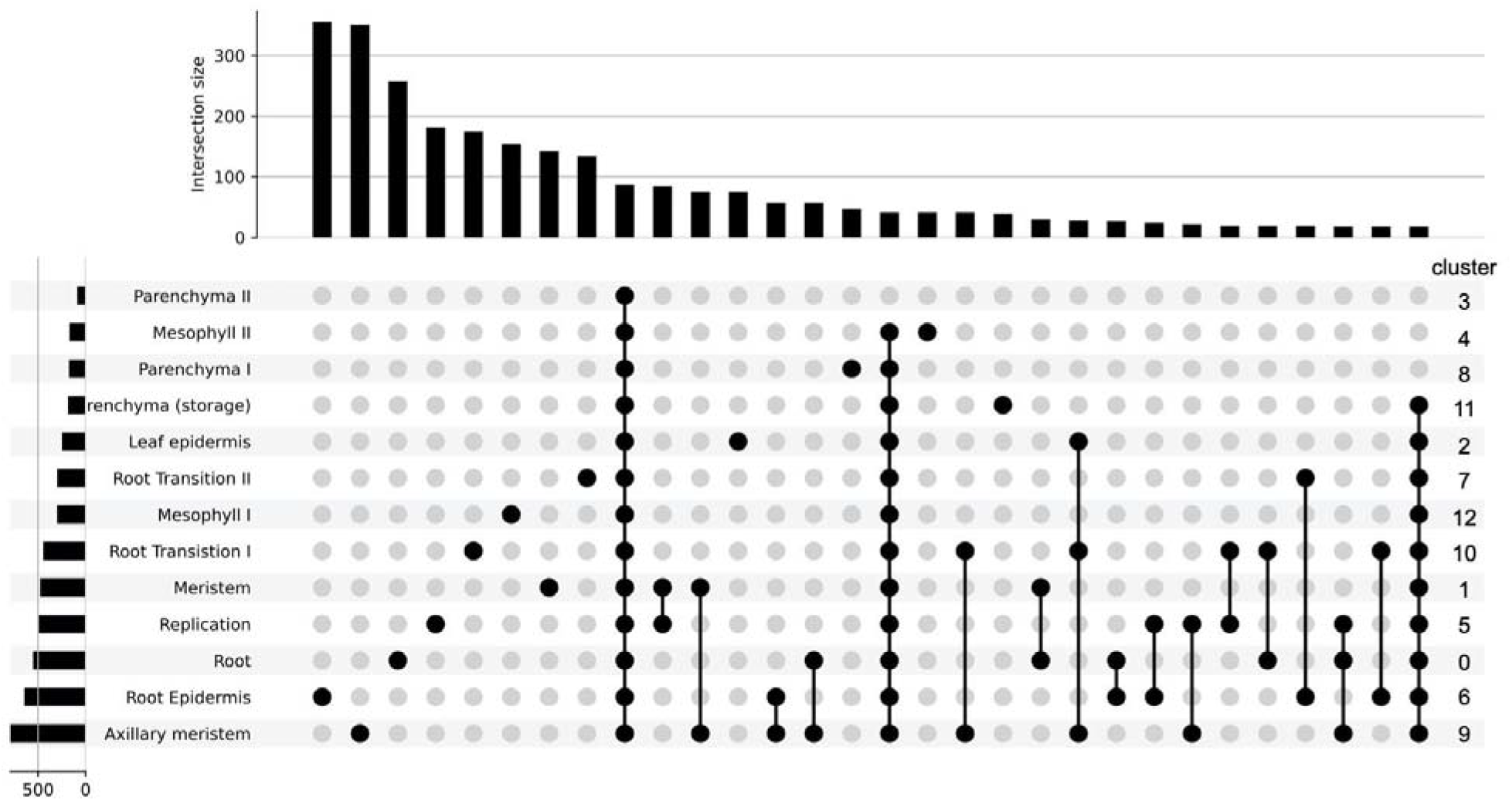
Cell type definitions supported by gene ontology (GO) categories. Upset plot showing the number of GO terms that are specific and overlapping across the thirteen different predicted cell types. The bars on top represent the number of intersecting GO terms per cell type, and the bars on the left represent the number of GO terms found in each cell type. The cell clusters as defined in Figure 5B are on the right hand side to highlight their correspondence with the predicted cell types.

**Figure 7.**
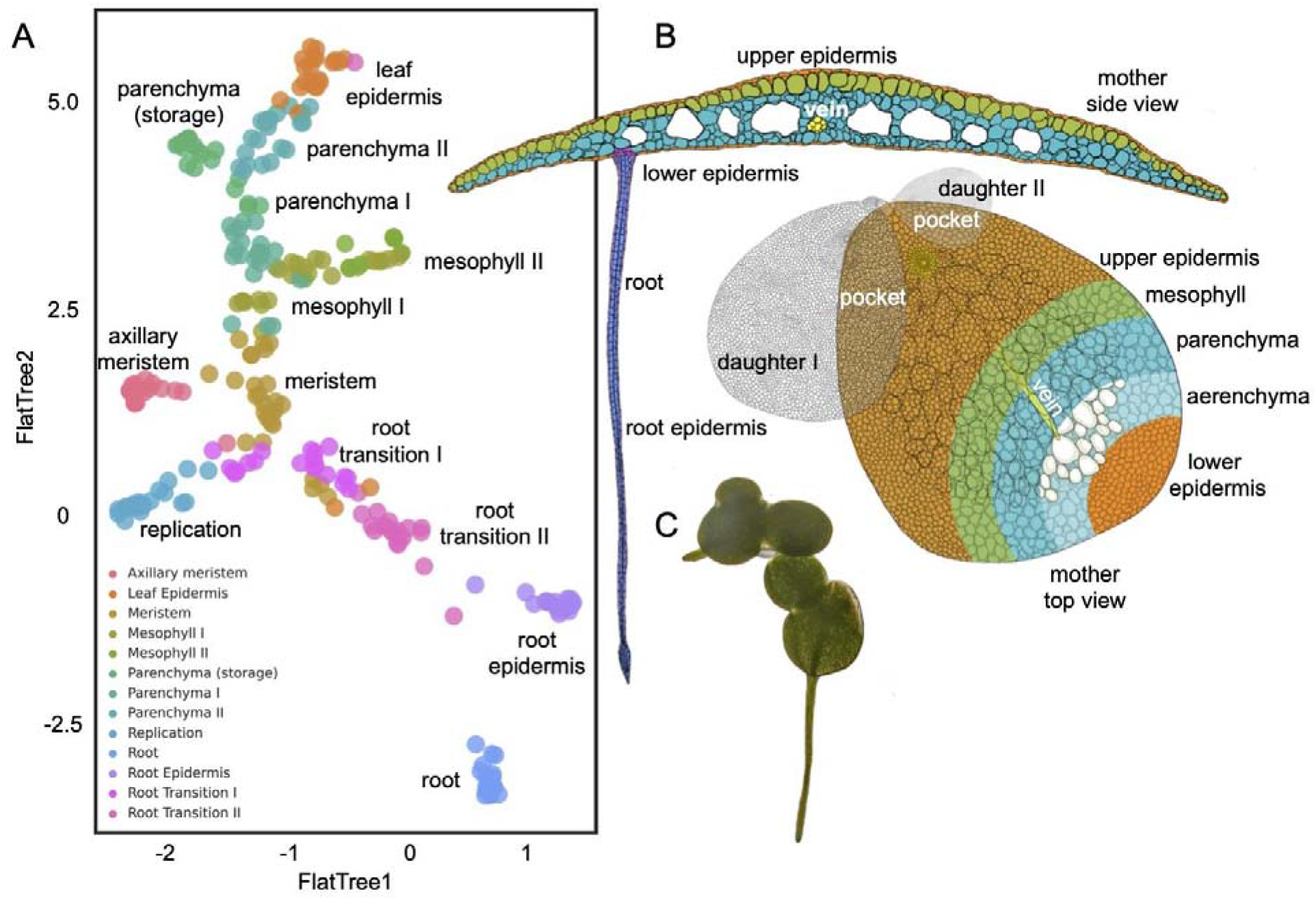
Trajectory analysis defines cell types of the entire *L. minuta* plant. A) UMAP tree embedding of snRNA-seq clusters describing the cell types. B) Cartoon of *L. minuta* based on (Banaszek and Musiał, 2011) coloring specific cell types per the UMAP. C) Live image of lm5633 showing the green roots.

### Meristematic-cell cluster

Following pseudo-time analysis, the gene expression profiles resulted in a large continuous cluster with multiple branches to terminally differentiated cell types (Fig. 5B and 7A). In the UMAP (Uniform Manifold Approximation and Projection) embedding we found several clusters relating to growth and division. Specifically, we defined the meristem (cluster 1) by an *TRANSLOCASE OF CHLOROPLAST 34 (TOC34), ACTIN RELATED PROTEIN (ARP4)* and the *ARGONAUT-like (AGO)* marker shown to be expressed in the shoot apical meristem in maize (Xu et al., 2021). A *TOC34* promoter GUS fusion in *A. thaliana* has been reported in the meristem of green tissues and root cells, and *ARP4* has been shown to function in flower development across species (Gutensohn et al., 2000; Pandey and Chaudhary, 2016). Furthermore, we surveyed the most representative GO terms for the meristem cluster and found that the top eight most representative terms relate to response to oxidative stress or removal of superoxide radicals, which could be associated with DNA damage protection (Supplemental Table S8). However, the representative set of GO terms relate to tissue development including GO:1905330, “regulation of morphogenesis of an epithelium;” GO:0090175, “regulation of establishment of planar polarity;” GO:2000023, “regulation of lateral root development;” GO:0048831, “regulation of shoot system development;” GO:0022603, “regulation of anatomical structure morphogenesis.” Finally, 12% of all GO descriptions related to development compared to terminally differentiated cell types such as “leaf epidermis” that contained only 3.6% of terms relating to development.

The placement of the meristem cluster in the UMAP embedding was supported by two adjacent clusters that we described as the “axillary meristem” and “replication” (or potentially endoreplication). The axillary meristem identification is supported by the presence of *LAX PANICLE1(LAX1)*, which is mediated at the transcript level in the axillary meristem (Oikawa and Kyozuka, 2009). There are also several genes involved in ribosome biosynthesis and genes coding for the ribosomal complex *(PRPL10, RLP24, RPA1A, RPL12-C, RPS16)*, which would suggest these cells are actively translating mRNA into proteins. The DNA replication/endoreduplication cell cluster contains *ETHYLENE INSENSITIVE (EIN2)* that leads to cell expansion through ethylene signaling and an *A-TYPE CYCLIN (CYCA3-1)*, which is critical for G1-to-S phase transition having a central role in the meristematic tissue (Takahashi et al., 2010; Feng et al., 2015). It is possible that these three specific clusters are all located in the meristematic-like tissue given their close proximity in the UMAP embedding and similarity in nuclei expression profiles. Both of these clusters share common GO terms, GO:0051093, “negative regulation of developmental process;” GO:0022603, “regulation of anatomical structure morphogenesis,” which suggest they are cells with defined processes similar to meristematic cells but different enough to warrant exclusive clustering.

The meristem cells give rise to all other cell types in clonally propagated plants, yet many of the transcription factors and networks leading to fated cell types are unknown. Transcription factor *WRKY32* has increased expression in the transition from meristem to green frond-like cell types, where it has been previously shown to be involved in ethylene signaling in tomato, where RNAi repression leads to yellowing of fruits (Zhao et al., 2021). It has been shown previously that related *L. minor* clones show reduced growth from exogenous ethylene treatment (Utami et al., 2018). Conversely, WRKY transcription factors associated with root cell transition are *WRKY6* and *WRKY65*. *WRKY6* is associated with pathogen defense, phosphate translocation, and arsenate resistance and *WRKY65* induces Jasmonate and salicylic acid responses relating to pathogen response (Huang et al., 2016; Wang et al., 2020). The expression of several WRKY transcription factors in the meristematic transition leading to terminal cell types suggests hormone signaling is a major contributing factor in lm5633 cellular development.

### Root-like tissues

The predicted meristem-like cells are centrally located between two branches leading to terminal cell types in the trajectory analysis (Fig. 5B and 7A). Although it has not been confirmed that duckweed roots are essential or act similarly to well developed root systems in model plants, we still define root cells by expression of two copies of the tandemly duplicated *PLANT LIPOXYGENASE 9* (*LOX9*) genes, which has been shown in soybean to be expressed in root nodules (Hayashi et al., 2008). Additional root makers are involved in metabolite transport by *HIGH-AFFINITY POTASSIUM TRANSPORTER 8* (*HAK8*) and *MANGANESE ATPASE TRANSPORTER (ECA2) (Mills et al., 2008)*. We find an additional marker gene *UBIQUITIN E2 CONJUGATING ENZYME* (*UBC13*) that has been shown to be involved in root development (Wen et al., 2014). We also find an additional paralog of *LOX9* in the root transition I cluster suggesting the paralogs are cell specific but concordant in their function progressing toward formation of root-like cells.

Visual morphologies have historically been used to define long-lived, stable cell types. Single cell/nuclei sequencing aims to capture all cell types including transitory cells. In some cases, snRNA-seq may capture transitional cell types that do not have well characterized visual properties or marker gene expressions but show progression to terminal cell types based on functional characterization of the snRNA-seq expression profile. Although difficult to define a specific cell type, here, the root transition II cluster contains a marker gene for *ASYMMETRIC LEAVES1* (*AS1*), which is essential for adaxial-abaxial leaf polarity and associated with an initial committed step towards root-like tissue development (Xu et al., 2003; Machida et al., 2015). Likewise, *SLEEPY1* (*SLY1*) is responsible for gibberellin signaling, cell growth and elongation, suggesting this cluster contains cells transitioning to the final root-like cells (McGinnis et al., 2003; Xu et al., 2003)*. ELONGATED HYPOCOTYL 5 (HY5)* is also found in this cluster and is known to play a role in induction of light induced genes and root gravitropism (Srivastava et al., 2015). It is interesting to note that lm5633 roots were exposed to light in this experiment since the plants are grown in clear flasks, which results in the roots appearing visually green and expressing light harvesting genes (Fig. 7C).

### Epidermal tissues

The exterior cells forming the epidermis provide a barrier from the rhizosphere and phyllosphere, and are often associated with increased production of hydrophobic waxes and cutins. Lipid biosynthesis is essential for wax and cutin production that provide extracellular protection in epidermal cells. Fatty acid biosynthesis,known to occur in epidermal cells,is mediated through the key marker genes *3-KETOACYL-COENZYME (KCS3)* and *LONG CHAIN ACYL-CoA SYNTHETASE (LACS2)(Kim et al., 2013)* (Supplemental Table S6). Additionally, root epidermal cells were defined based on expression of *CRINKLY 4* (*CR4*), which is known to have a role in maize epidermal cell formation (Becraft et al., 1996). Interestingly, frond epidermal cells are located in close proximity to root epidermal cells in UMAP embedding consistent with these cell types sharing similar expression despite having different cell trajectories (Fig. 5B and 7). The frond epidermal cells were defined by the marker gene *ECERIFERUM-like*, which has been shown to be important for cuticle wax development (Haslam et al., 2017). Additional markers include an auxin efflux carrier in starch metabolising cells *PIN-FORMED (PIN7)* aiding root termination (Kim et al., 2013; Rosquete and Kleine-Vehn, 2018). This is consistent with this cluster’s position in the UMAP embedding (Fig. 5B) in close proximity to the root epidermis but highly diverged in the complement of expressed photosynthetic genes and complete divergence in the trajectory analysis (Fig. 7A). Likewise, this cell type contains genes *ATP-DEPENDENT ZINC METALLOPROTEASE (FTSH2)*, two orthologues of *DEFECTIVE KERNEL 1 (DEK1)* and *CYTOCHROME C OXIDASE 15 (COX15)*. These genes are involved in carbon storage where *FTSH2* has been shown to be involved in thylakoid biosynthesis and *COX15* shows increased expression in response to decreased cellular respiration (Vishwakarma et al., 2015; Haslam et al., 2017). Similarly, DEK1 is expressed in aleurone-like cells in maize that are involved in starch metabolism (Tian et al., 2007). It has been shown that *DEK1* also has adverse effects on leaf morphology and it is likely that this cluster contains frond epidermal cells.

### Frond tissues

Mesophyll, located in leaf or frond tissue, are the primary cells involved in photosynthetic light and CO_2_ capture for the generation of sugars to sustain the plant. Mesophyll-like cells were determined based on significant expression of multiple marker genes associated with light perception, thylakoid biogenesis, and cytokinin degradation related to stress by *YELLOW STRIPE-LIKE (YSL9)* and *THYLAKOID LUMEN PPIase* (*TLP40*). *YSL9* in rice is associated with leaf vasculature but in this case is potentially more similar to mesophyll like cells in this basal aquatic monocot (Wen et al., 2020). *TLP40* has been shown to be expressed in mesophyll and at 2-fold greater expression in bundle sheath cells (Vojta et al., 2019). Interestingly, the representative GO terms suggest these cells are carbon limited and the photorespiratory cycle is engaged where the top GO terms are mitochondrial transport of glycolate (GO:0006626, “protein targeting to mitochondrion;” GO:0072655, “establishment of protein localization to mitochondrion;” GO:1901975, “glycerate transmembrane transport;” GO:0097339, “glycolate transmembrane transport”).

Parenchyma cells in any given tissue can become specialized for a variety of purposes, yet in lm5633 it appears the specialization of parenchyma is geared toward photosynthetic metabolism and storage of photosynthate. Parenchyma I and II clusters are defined *EXORDIUM-like (EXO), SUCROSE SYNTHETASE 1 (SUS1), STARCH EXCESS 4 (SEX4), XAP5 CIRCADIAN TIMEKEEPER (XCT)* which promotes ethylene response in aerial tissue (Ellison et al., 2011). *SUS1* is a prominent player in the accumulation of terminally synthesized photosynthate in source tissues and *SEX4* mutants in leaf tissues promote starch metabolism (Kötting et al., 2009; Ma et al., 2014; Shi et al., 2019). Mutants of *EXO* have decreased epidermis, palisade and spongy parenchyma in *A. thaliana* and suggests *EXO* may be inhibiting differentiation to a terminal cell type (Schröder et al., 2011; Florian Schröder, Janina Lisso, Peggy Lange & Carsten Müssig). Marker genes in these clusters point to photosynthetic metabolism. A terminal cluster is observed branching off the parenchyma-like cells (Fig. 7A) denoted here as “parenchyma (storage)” since cells are expressing two copies of *ALPHA AMYLASE 2 (AMY2)* and *IMBIBITION INDUCIBLE (IMB1)*. The combination of these marker genes in a cell type may mean these cells are actively metabolizing starch in the vacuoles.

### Specific cell expression associated with invasiveness

*Lemna minuta* can occupy a variety of ecosystems due to their adaptability in absorption and tolerance of excess micro- and macro-nutrients. The uptake and adaptability of *L. minuta* has led to its use as a wastewater detoxifying/phytoremediation species (Fernández et al., 2020). Wastewaters are often contaminated with a variety of toxic compounds yet the exact mechanisms of plants ability to adapt to these compounds, including high levels of essential elements and heavy metals, remains unclear (Frick, 1985; Davis et al., 2002; Del-Campo Marín and Oron, 2007; Gür et al., 2016; Liu et al., 2018; Türker et al., 2019). We find increased expression of *BORON TRANSPORTER 4* (*BOR4*) in the mesophyll and replicating cells (Fig. 8). We also found a large TD (11 copies; eighth largest) of a boric acid transporter (MIP aquaporin) (Fig. 3C; Supplemental Table S4). We find similar increased expression of *IRON-REGULATED TRANSPORTER 2* (*IRT2*) and *METAL-NICOTIANAMINE TRANSPORTER 9 (YSL9)* in the mesophyll cells. *IRT2* also shows a TD expansion (4 copies with 2 copies expressed here), although modest compared to *BOR4*. Overaccumulation of boron and toxic heavy metals can have an adverse effect on plant growth yet *L. minuta* grows and accumulates heavy metals readily (Gür et al., 2016). This is compelling evidence that *L. minuta’s* invasiveness and adaptability in wastewater may be, at least in part, due to the increased *BOR4* expression, as *BOR4* is essential for boron export from cells. Further snRNA-seq under micro- and macro-nutrient stress conditions would greatly improve our understanding of cell specific responses and adaptation to abiotic stressors.

**Figure 8.**
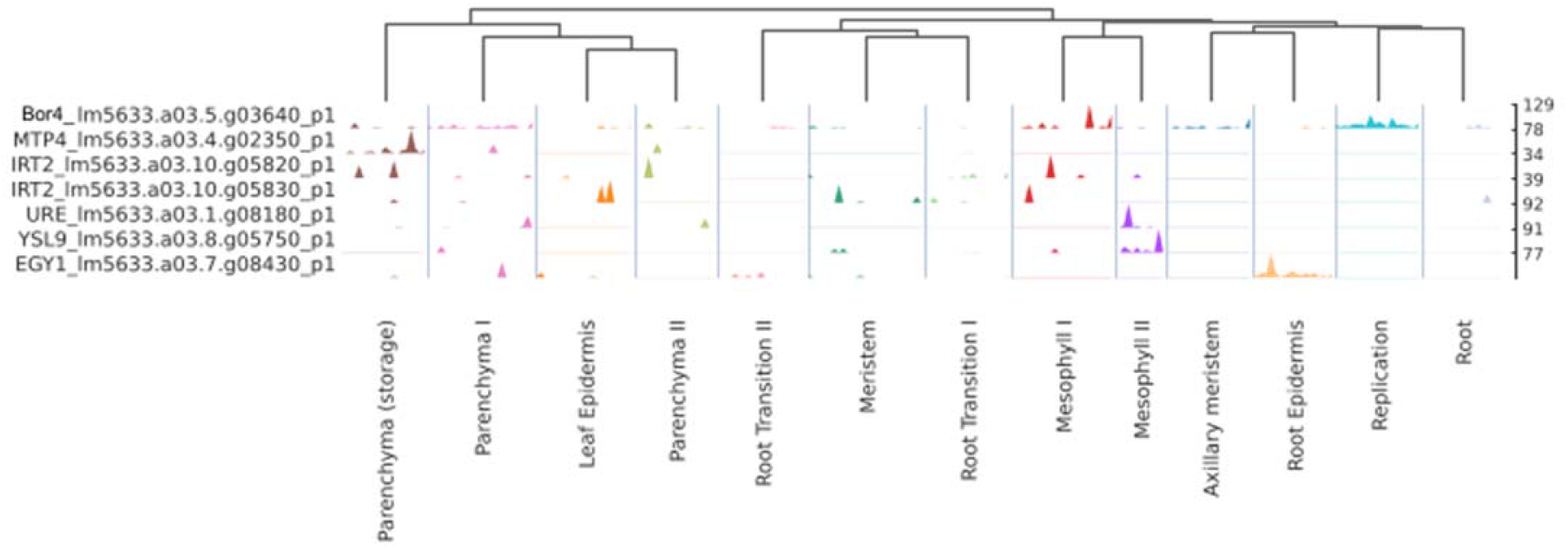
Metal transport and accumulation associated genes in *L. minuta* are expressed in mesophyll cells. SnRNA-seq expression profiling of metal tolerance related genes reveal a cell type specific expression pattern with greater gene expression localized to the mesophyll cell types in lm5633. Lm5633.a03.5.g03640 (top) encodes *BORON TRANSPORTER 4 (BOR4)* localized to mesophyll and replicating cells described further in the text.

## Discussion

Here we endeavoured to define the trajectory of cells in a morphologically reduced, fast growing non-model plant useful for phytoremediation but also invasive in specific environments. We generated a chromosome-resolved genome for the Lesser Duckweed *L. minuta* clone lm5633 and found that it did not share a lineage specific WGD with *S. polyrhiza* yet had an increased number of Tandem Duplications (TDs) for genes involved in pathogen defence, response to stress and nutrient acquisition. Additionally, we performed snRNA-seq on a clonal population of whole *L. minor* plants that represent all developmental stages, which enabled us to discern differentiated cell types and their development from the meristematic cells. Finally, the data provided evidence that the mesophyll is a site of elemental and heavy metal transporter expression.

Duckweed are particularly suited for single cell studies due to their size, direct contact with the media, small non-redundant genome, and rapid clonal growth (Fig. 1). Most single cell studies to date have been conducted using protoplasting on root or reproductive structures, which have the advantage of having all cells in the developmental continuum (Denyer et al., 2019; Ryu et al., 2019; Liu et al., 2020a). Similarly, Duckweed provide a continuum of all cell types and developmental stages due to their rapid clonal growth and nested generations. Moreover, Duckweed are in direct contact with their media allowing for detailed and controlled manipulation of the environment, which in the the past was leveraged to understand auxin biology (Slovin and Cohen, 1988). The snRNA-seq method described here also couples well with experiments where abiotic/biotic treatments of the samples are desired because they can be immediately frozen and transcriptomic perturbations measured additionally frozen nuclei presumably capture actively transcribing mRNA. One potential downside of snRNA-seq is the loss of RNA in the cytoplasm; more experiments comparing the different techniques will be required to evaluate this limitation.

Additionally, the Duckweed genome provides a compelling platform for gene annotation and functional analysis. It was shown that *S. polyrhiza* has a reduced set of non-redundant protein coding genes even though it has had two lineage specific WGD events (Wang et al., 2014). One challenge in model species and crops is the number of paralogs (family size) that complicate gene functional analysis due to redundancy of action. Here, the lm5633 genome also has a streamlined set of genes that shows a similar reduction in paralogs compared to other plants (Fig. 4A). However, lm5633 does have a much higher level of fractionation compared to sp9509 where lm5633 has 4.1% retention of syntenic paralogs compared to 49% retained in sp9509, which means that either lm5633 is evolving at a greater rate or it has a different WGD history. We find evidence that both lm5633 ands wa8730 lack the *S. polyrhiza* lineage specific WGD and only have the shared monocot *tau* WGD (Fig. 4, B and C). Currently there are conflicting reports whether *S. polyrhiza* has the *tau* WGD (Wang et al., 2014; Ming et al., 2015; Hasing et al., 2020). One explanation is that the lineage-specific WGD has obscured the *tau* WGD in *S. polyrhiza*, and that lm5633 is more fractionated due to the very ancient (~150 mya) WGD. More Duckweed genomes will help resolve the WGD history.

All single cell studies are currently in model systems, yet as we have shown here one can go from a wild plant to a chromosome-resolved genome and leverage snRNA-seq dataset to generate functional information. While model plants and specifically *A. thaliana* have high quality annotations and marker genes for cells, there are significant challenges in defining marker genes or assigning cellular functions for even well studied plants and crops. Here we leveraged a three tiered approach to assign cell type. Our realization was that cell trajectory analysis provides rich information about cell type, especially when it is coupled with a developmental series shown here using an entire Duckweed plant similarly in roots and floral tissues (Denyer et al., 2019; Satterlee et al., 2020; Xu et al., 2021). GO and orthology comparisons are more fraught with issues due to sparse datasets (i.e. few or no GO terms associated with gene predictions) where orthology may not exist or be cryptic with model species or crops. However, when these three criteria are leveraged, annotation can be improved as well as provide support for known and new cell types, like transition cell types.

The trajectory of cell types from meristem to terminally differentiated cells provides clues as to their function. Cellular studies have suggested that the meristem is unique in Duckweed since it gives rise to daughter fronds (Landolt, 1986). We found the meristem is divided into three potential cell clusters (meristem, replication, and axillary meristem), which provides a clue as to how duckweed is dividing rapidly leading to continuous daughter frond production. The overlapping trajectory and GO terms of mersitem-like and root-like cells (Fig. 6 and 7), suggests the root may play additional roles that are unknown at this time, and may clarify its usefulness. Furthermore, root and frond epidermal cells have distinct trajectories (Fig. 7) yet arrive in a similar expression space consistent with their overall cellular function (Fig. 5B). Finally, the emergence of the photosynthetic mesophyll-like cells from the meristem then giving rise to the metabolically active and terminally differentiated parenchyma cells. We did not identify stomatal cells or vascular tissue (vein) cells and this could be a result of our methodology or the fact that *L. minuta* only has one vein and about 30 stomata (~60 cells per plant) (Landolt, 1986). Since each mother frond has 256 (each daughter frond has 2 or more generations) developing, yet unexpanded daughter/grand-daughter fronds the majority of the cells we detect will be actively dividing; deeper sequencing will be required to accurately identify small populations of terminally differentiated cells.

In model plants, nutrient uptake and transport primarily occurs in the roots and vasculature (Yan et al., 2020). However, the Duckweed frond is in direct contact with its environment, which may explain why we detect high expression of nutrient uptake genes in the cells we define as the photosynthetically active mesophyll (Fig. 8). Both *ITR2* and *BOR4*, which are also both expanded in lm5633 through TDs, are highly expressed in mesophyll. However, it has been shown that *IRT2* is expressed in the roots (Vert et al., 2001), and *BOR4* is expressed in the xylem in *A. thaliana* (Takano et al., 2002). The solitary vein of lm5633 is in close proximity to the mesophyll, which makes it formally possible that we have mis-identified some of these cells and they actually are vasculature or at an expression level these cells are too similar to separate with the current sequencing coverage. Either way, this simplified system provides an opportunity to further define the acquisition, transport and storage of toxic compounds at a single cell level. Future studies will enable clarification of cell types as well as provide the opportunity to dissect the specific cellular mechanisms associated with phytoremediation and invasiveness.

### Conclusions

Duckweed provides an unprecedented opportunity to study the cell fates across an entire morphologically reduced plant. Coupled to snRNA-seq, this system provides a new opportunity to study the cell-specific responses to environmental changes. This dataset represents a first look and will surely be refined with additional datasets in additional plants. Different plants, conditions and treatments will refine our understanding of cell type and ultimately identify new cell types and niches.

## Materials and Methods

### Plant collection and growth

*Lemna minuta* was collected on October 10, 2019 from Cotton Creek Park, Encinitas, CA, USA (33°2′58″N 117°17′29″W), which is a waste water slough close to the ocean (Fig. 1A). A population of clones were collected, surface sterilized and one clone was retained to multiply the population. The representative clone was deposited in the Rutgers Duckweed Stock Collective (RDSC; www.ruduckweed.org) and was assigned the clone number lm5633. Plants were propagated on Schenk and Hildebrandt (SH) media as described at RDSC.

### Genome sequencing

High Molecular weight (HMW) DNA was extracted with modifications (Lutz et al., 2011). The resulting DNA was quality controlled for size and contamination. Unsheared HMW DNA was used to make ONT ligation-based libraries. Libraries were prepared starting with 1.5 ug of DNA and following all other steps in ONT’s SQK-LSK109 protocol. Final libraries were loaded on a ONT flow cell (v9.4.1) and run on the GridION. Bases were called in real-time on the GridION using the flip-flop version of Guppy (v3.1). The resulting fastq files were concatenated and used for downstream genome assembly steps. Illumina 2×150 paired end reads were also generated for genome size estimates and polishing genome sequences. Libraries were prepared from HMW DNA using NEBNext Ultra II (NEB, Beverly, MA) and sequenced on the Illumina NovaSeq. Resulting raw sequence was only trimmed for adaptors, resulting in >60x coverage.

### Genome size estimation

K-mer (k=31) frequency was estimated with Illumina paired end reads (2×150 bp) using Jellyfish (v2.3.0) (Marçais and Kingsford, 2011) and analyzed using in house scripts and GenomeScope and GenomeScope2 (Vurture et al., 2017; Ranallo-Benavidez et al., 2020). Genome size, heterozygosity and repeat content were first estimated with GenomeScope (http://qb.cshl.edu/genomescope) (Table 1). The K-mer frequency plot is consistent with a highly heterozygous diploid genome or a tetraploid genome; based on previous flow cytometry, *L. minuta* is diploid with an average genome size of 365 Mb (Table 1) (Bog et al., 2020).

### Genome assembly

Resulting fastq files passing QC were assembled using our previously described pipeline (Michael et al., 2018) with the modification that the initial assembly was generated using FlyE (Kolmogorov et al., 2019). The resulting assembly graph (gfa) was visually inspected with Bandage (v0.8.1) (Wick et al., 2015), which revealed a branching pattern consistent with a heterozygous genome with structural differences between haplotypes. Consensus was generated with three (3) iterative cycles of mapping the ONT reads back to the assembly with minimap2 followed by Racon (v1.3.1) (Vaser et al., 2017), and the final assembly was polished iteratively three times (3) using 2×150 bp paired-end Illumina reads mapped using minimap2 (v2.17-r941) (>98% mapping) followed by pilon (v1.22) (Walker et al., 2014). The resulting assembly was assessed for traditional genome statistics including assessing genome completeness with Benchmarking Universal Single-Copy Orthologs (BUSCO) (v3) liliopsida odb10 database (Table 1) (Simão et al., 2015).

### High throughput Chromatin Conformation Capture (HiC) genome scaffolding

Crosslinking was performed on ground tissue and nuclei were isolated following (Colt). HiC data was generated using the Arima-HiC Kit User Guide for Plant Tissues (Link et al., 2018), according to the manufacturer’s protocols (Catalog #A510008 Document Part #A160135 V00). Libraries were generated following the Arima-HiC Kit Library Preparation Guide for Swift Biosciences Accel-NGS 2S Plus DNA Library Kit (Catalog #A510008 Document Part #A160140 V00) and sequenced on Illumina NovaSeq. We used standard methods defined in the Aiden lab genome assembly cookbook (https://github.com/aidenlab/3d-dna/). The primary steps include alignment of HiC reads to the polished lm5633 assembly and creation of a contact map using the Juicer pipeline (https://github.com/aidenlab/juicer) followed by automated scaffolding using 3d-dna. The scaffolds were then inspected and manually corrected with Juicebox Assembly Tools (JBAT) before being finalized by the 3d-dna post review pipeline. The resultant scaffolds for lm5633 were output and renamed from longest to shortest.

### High-copy repeat analysis

Long read ONT assemblies provide another measure of completeness through the identification of high copy repeats such as centromeres and telomeres sequences (VanBuren et al., 2015). We employed a searching strategy to identify the centromeres that leverages the idea that the highest copy number tandem repeat (TR) will be the centromere in most genomes (Melters et al., 2013). We searched the genomes using tandem repeat finder (TRF; v4.09) using modified settings (1 1 2 80 5 200 2000 -d -h) (Benson, 1999). TR were reformatted, summed and plotted to find the highest copy number TR per our previous methods (VanBuren et al., 2015). While lm5633 had robust telomere arrays (Table 1), we could not detect a high-copy number centromere repeat similar to what we have found in *S. polyrhiza* (Michael et al., 2017), which could mean lm5633 has holocentric centromeres.

### Gene prediction and annotation

The chromosome resolved lm5633 genome was annotated using a pipeline consisting of four major steps: repeat masking, transcript assembly, gene model prediction, and functional annotation. Repeats were identified using EDTA (v1.9.8) (Ou et al., 2019) and these repeats were used for softmasking. ONT cDNA reads were aligned to the genomes using minimap2 and assembled into transcript models using Stringtie (v1.3.6). Softmasked genomes and Stringtie models were then processed by Funannotate (v1.6) (https://github.com/nextgenusfs/funannotate) to produce gene models. Predicted proteins were then functionally annotated using Eggnog-mapper (v2) (Huerta-Cepas et al., 2017). The resulting gene models were renamed reflecting the chromosome and the linear position on the chromosome.

### Orthogroup analysis and synteny

Gene families and overrepresented groups were determined with Orthofinder (v2.4.0) and CAFE5 (https://github.com/hahnlab/CAFE5). Genomes were accessed from Phytozome13 (https://phytozome-next.jgi.doe.gov/) or from specific publications such as the *Colocasia esculenta* (Taro) (Yin et al., 2021), *Spirodela intermedia* (Hoang et al., 2020), *Spirodela polyrhiza clone 9509* (Hoang et al., 2018) and *Wolffia australiana* clone 8730 (Michael et al., 2020). Orthofinder results were parsed and used for CAFE5 by modifying the species tree with “make_ultrametric.py” and filtering orthocounts.tsv with “clade_and_size_filter.py”. Gene trees were visualized with iTOL (Letunic and Bork, 2019). The SynMap tool on CoGe (Grover et al., 2017) and McScan python version (https://github.com/tanghaibao/jcvi/wiki/MCscan) were utilized to generate whole genome synthetic maps, identify syntenic orthologs, estimate synonymous substitutions (Ks) across genomes, and generate figures.

### *Single nuclei RNAseq* (snRNA-seq)

We generated a snRNA-seq dataset from a population of whole lm5633 plants grown in 250 ml erlenmeyer flasks under 12 hours of light and 12 hours of dark (intermediate days) with constant 22°C temperature (LDHH). SnRNA-seq was performed using the SMARTseq2 protocol on nuclei isolated from frozen plant material as previously described (Bakken et al., 2018) with some modifications. Nuclei were extracted with a custom buffer consisting of Tris-HCL (9.5) 15mM, EDTA 10mM, KCl 130mM, NaCl 20mM, PVP-10 8%, Spermine 0.07g, Spermidine 0.05g, Triton X-100 0.10%, BME 7.50%. Individual nuclei were sorted on the FACsAria Fusion system into SMARTseq2 lysis buffers allocated in 96 wells. The resulting libraries were sequenced on the Illumina NovaSeq platform. The following QC was performed on the resulting reads: genes were filtered if they were expressed in fewer than 5 cells, cells were required to have at least 100 genes expressed and at least 20,000 reads mapped to the transcriptome; this QC resulted in 269 cells with 8,457 genes expressed. An expression matrix was generated for each cell by each gene. We leveraged Single-cell Trajectories Reconstruction, Exploration And Mapping (STREAM) to cluster and map the cell trajectories of the 269 sequenced nuclei (Chen et al., 2019). Clustering was performed with K-means (k=13) in two components and 26 nearest neighbors using the top 100 principal components based on maximized silhouette scores. A weighted centroid principle graph was generated with epg_alpha=0.01,epg_mu=0.05,epg_lambda=0.05,jobs=10 and refined with st.extend_elastic_principal_graph using epg_ext_par=0.8. To detect significant marker genes st.detect_markers with cutoff_zscore=1.0 and cutoff_pvalue=0.01 was used. Transition markers were determined with st.detect_transition_markers using cutoff_spearman=0.4 and cutoff_logfc=0.25. SCANPY was used for data visualization purposes (Wolf et al., 2018).

The 269 nuclei formed 13 clusters that we defined using gene ontology (GO), KEGG, and PFAM annotation into cell types based on orthologous marker genes. Orthology was determined with Orthofinder (Emms and Kelly, 2019) as above. Marker genes from model species based on existing literature were used to search the orthogroups for lm5633 orthologues (Supplemental Table S6). The marker genes in model organisms were compared with the orthofinder gene table and a corresponding lm5633 ortholog was assigned. Generally, annotations with a geneID from eggnog mapper were more reliable than observing marker genes in larger gene families (>3 copies per genomes). GO terms and quantities for cluster marker genes were parsed and form marker genes lists using in-house scripts. GO lists were visualized using REVIGO and CirGO (Supek et al., 2011; Kuznetsova et al., 2019).

## Author Contributions

TPM and BWA conceived the study, BWA and MN carried out the lab work, KC performed sequencing, BWA, BDA, NH and TPM performed the analysis, RHS and TPM oversaw the project, and TPM and BWA wrote the manuscript.

## Accession numbers

The final genome assembly is available on CoGe (https://genomevolution.org/) under the ID:61245 or Biosample: SAMN19243672. Genomic and snRNA-seq reads can be found in SRA under Biosample: SAMN19243672.

## Large datasets

Supplemental Table S1. BUSCO scores for lm5633.

Supplemental Table S2. Predicted repeat types in the lm5633, sp9509 and wa8730 genomes.

Supplemental Table S3. A comparison of telomere sequence lengths in Duckweeds.

Supplemental Table S4. Tandem duplications (TDs) in lm5633.

Supplemental Table S5. Previously published marker genes used for defining snRNA-seq cell types.

Supplemental Table S6. Orthogroups across all species.

Supplemental Table S7. Lemna minuta (lm5633) snRNA-seq marker genes per cell type.

Supplemental Table S8. Gene ontology (GO) terms for the meristem cluster.

## Acknowledgements

We thank Rutgers Duckweed Stock Cooperative http://www.ruduckweed.org/ for hosting the live duckweed strain under lm5633.

